# Novel proteostasis reporter mouse reveals different effects of cytoplasmic and nuclear aggregates on protein quality control in neurons

**DOI:** 10.1101/2020.11.09.374231

**Authors:** Sonja Blumenstock, Elena Katharina Schulz-Trieglaff, Anna-Lena Bolender, Kerstin Voelkl, Paul Lapios, Jana Lindner, Mark S. Hipp, F. Ulrich Hartl, Rüdiger Klein, Irina Dudanova

## Abstract

The cellular protein quality control machinery is important for preventing protein misfolding and aggregation, and decline in protein homeostasis (proteostasis) is believed to play a crucial role in age-related neurodegenerative disorders. However, how proteostasis capacity of neurons changes in different diseases is not yet sufficiently understood, and progress in this area has been hampered by the lack of tools to monitor proteostasis in mammalian models. Here, we have developed reporter mice for *in vivo* analysis of neuronal proteostasis. The mice express EGFP-fused firefly luciferase (Fluc), a conformationally unstable protein that requires chaperones for proper folding and sensitively reacts to proteotoxic stress by formation of intracellular Fluc-EGFP foci and by reduced luciferase activity. Using these mice, we provide evidence for proteostasis decline in the aging brain. Moreover, we find a marked impairment in proteostasis in tauopathy mice, but not in Huntington’s disease mice. Mechanistic investigations in primary neuronal cultures demonstrate that cytoplasmic, but not nuclear, aggregates cause defects of cellular protein quality control. Thus, the Fluc-EGFP reporter mice enable new insights into proteostasis alterations in different diseases.

## Introduction

Maintaining the integrity of the cellular proteome is essential for survival. The cellular protein quality control system safeguards protein homeostasis (proteostasis) by ensuring correct folding of new proteins, detecting and refolding damaged proteins, and targeting terminally misfolded proteins for degradation (Balchin et al., 2016; Klaips et al., 2018). Age-dependent decline in protein quality control is believed to play a crucial role in neurodegenerative diseases, a group of brain disorders characterized by aggregation of misfolded proteins and neuronal cell death, such as Alzheimer’s, Parkinson’s and Huntington’s disease (HD) (Soto and Pritzkow, 2018). Enhancing the capacity of the protein quality control system has therefore emerged as a promising therapeutic strategy for neurodegenerative proteinopathies (Klaips et al., 2018; Smith et al., 2015). However, our current knowledge about the proteostasis changes *in vivo* during disease progression is still scarce. While attempts to ameliorate aggregate toxicity by upregulating chaperones have been successful in cell culture, fly and worm models (Auluck et al., 2002; Carmichael et al., 2000; Hageman et al., 2010; Kuo et al., 2013; Outeiro et al., 2006; Vos et al., 2010; Wu et al., 2010), they have produced less satisfactory results in mammalian models (Hansson et al., 2003; Krishnan et al., 2008; Labbadia et al., 2012; Liu et al., 2005; Sharp et al., 2008; Shimshek et al., 2010; Xu et al., 2015; Zourlidou et al., 2007). Reliable genetic reporters that allow monitoring the status of cellular proteostasis *in vivo* are essential for understanding disease mechanisms and for assessing the efficacy of potential treatments targeting the protein quality control system. Thus, transgenic mice expressing ubiquitin-proteasome system (UPS) reporters have been used successfully for investigating protein degradation in disease models (Bett et al., 2009; Cheroni et al., 2009; Kristiansen et al., 2007; Lindsten et al., 2003; Myeku et al., 2016; Ortega et al., 2010). However, tools for monitoring proteostasis in general are still lacking.

Wildtype and mutated versions of the conformationally unstable firefly luciferase (Fluc) protein fused to EGFP are ideal for use as proteostasis sensors and have proven valuable in cell lines and in *C. elegans*(Donnelly et al., 2014; Gupta et al., 2011). These sensors depend on cellular chaperones for proper folding and enzymatic activity. Proteotoxic conditions that overload the protein quality control system lead to misfolding of Fluc-EGFP, which can be revealed by two readouts: decrease in bioluminescence due to decline in luciferase activity, and formation of Fluc-EGFP foci in the cell as a result of decreased solubility of the misfolded sensor (Gupta et al., 2011).

To gain a deeper understanding of proteostasis changes in mouse disease models, we generated new reporter mice expressing Fluc-EGFP in the nervous system. Using these mice, we reveal unexpected differences in proteostasis caused by protein aggregates located in different cellular compartments.

## Results

### Fluc-EGFP sensor reacts to proteostasis changes in primary neurons

We first asked whether Fluc-EGFP variants can be used as reporters of proteotoxic stress in primary neurons. In the following experiments we used two versions of Fluc, wildtype (FlucWT) as well as single mutant FlucR188Q (FlucSM, both with a C-terminal HA tag). FlucSM is a conformationally destabilized mutant that was previously shown to have higher sensitivity to proteotoxic stress than FlucWT (Gupta et al., 2011), however its expression levels in neurons were relatively low, possibly due to efficient degradation. Transfection of Fluc-EGFP constructs did not lead to toxicity in murine primary cortical cultures, as demonstrated by immunostaining against the apoptotic marker cleaved Caspase-3 (Fig. S1A-B). Cultures transfected with Fluc-EGFP constructs were then subjected to several treatments to induce proteotoxic stress. MG-132 (5 μM for 4 hours) was used to inhibit protein degradation by the proteasome, Bafilomycin A1 (10 nM for 24 hours) was used to inhibit autophagy, and 17-AAG (0.5 μM for 4 hours) was used to inhibit Hsp90, a major cytosolic chaperone. All these treatments induced a change in the distribution of Fluc-EGFP, which typically formed several small compact foci in the perinuclear region, while the intensity of diffuse EGFP fluorescence in the rest of the cytoplasm decreased (Fig. 1A-B and S1C). In addition, cells were subjected to heat shock at 43°C for 30 min. This treatment led to an even stronger Fluc-EGFP response, with many cells showing a complete loss of diffuse cytoplasmic EGFP fluorescence and formation of multiple Fluc-EGFP foci throughout the cytoplasm (Fig. 1A-B and S1C).

**Figure 1.**
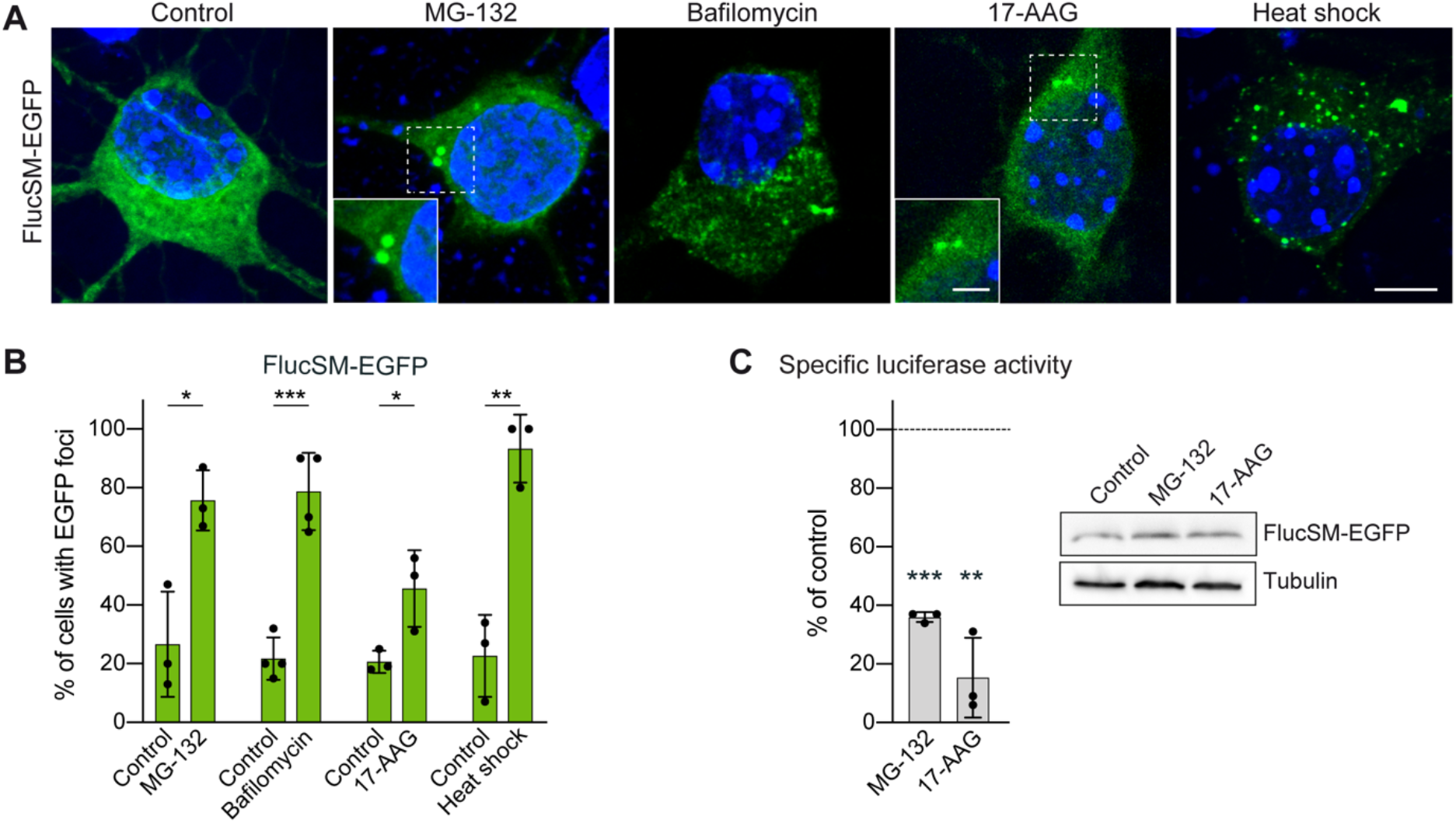
Fluc-EGFP reacts to proteostasis changes in primary neurons. **A**, Representative images of DIV 3+2 cortical neurons transfected with FlucSM-EGFP (green) and subjected to the indicated treatments. Nuclei are labeled with DAPI (blue). Insets show higher magnification of the areas outlined by dashed boxes. **B**, Quantification of FlucSM-EGFP foci formation in transfected neurons. N=3 independent experiments, 40-45 neurons/condition/experiment; Two-tailed t-test. **C**, Left, quantification of specific luciferase activity of FlucSM-EGFP upon indicated treatments, normalized to respective vehicle-treated controls. N=3 independent experiments; One-sample t-test. Right, representative Western blots of neuronal lysates for the indicated conditions. Tubulin was used as a loading control. *p<0.05, **p<0.01, ***p<0.001. Scale bars: A, 5 μm; insets, 2 μm.

We next used the luciferase assay to evaluate the enzymatic activity of Fluc-EGFP as an additional readout of proteostasis alterations. Luciferase activity measurements were normalized to Fluc-EGFP protein quantity determined by Western blot to obtain specific activity values. As expected, proteasome inhibition with MG-132 and Hsp90 inhibition with 17-AAG both resulted in a significant decrease in specific luciferase activity of FlucSM-EGFP by ~65% and ~85%, respectively (Fig. 1C). Overall, the response of Fluc-EGFP to heat shock and small-molecule inhibitors in primary neurons appeared stronger than the response observed in non-neuronal cell lines (Gupta et al., 2011), probably due to the high sensitivity of neurons to proteotoxic stress. Taken together, these results demonstrate that Fluc-EGFP can be reliably used to detect proteostasis disturbances in primary neurons.

### Fluc-EGFP reporter mouse for *in vivo* analysis of proteostasis

For *in vivo* studies of the protein quality control system in mouse models, we generated transgenic mouse lines expressing FlucWT-EGFP or FlucSM-EGFP under the control of the prion protein (PrP) promoter (Fig. 2A). In line with our observations in primary neurons, FlucSM-EGFP mouse lines showed rather low expression of the sensor. For further experiments, we selected the FlucWT-EGFP line 1214 (from here on, Fluc-EGFP mice), which had a broad salt-and-pepper-like expression in neurons throughout the brain, including regions affected in neurodegenerative proteinopathies such as the neocortex, hippocampus, basal ganglia and cerebellum (Fig. 2A-B).

**Figure 2.**
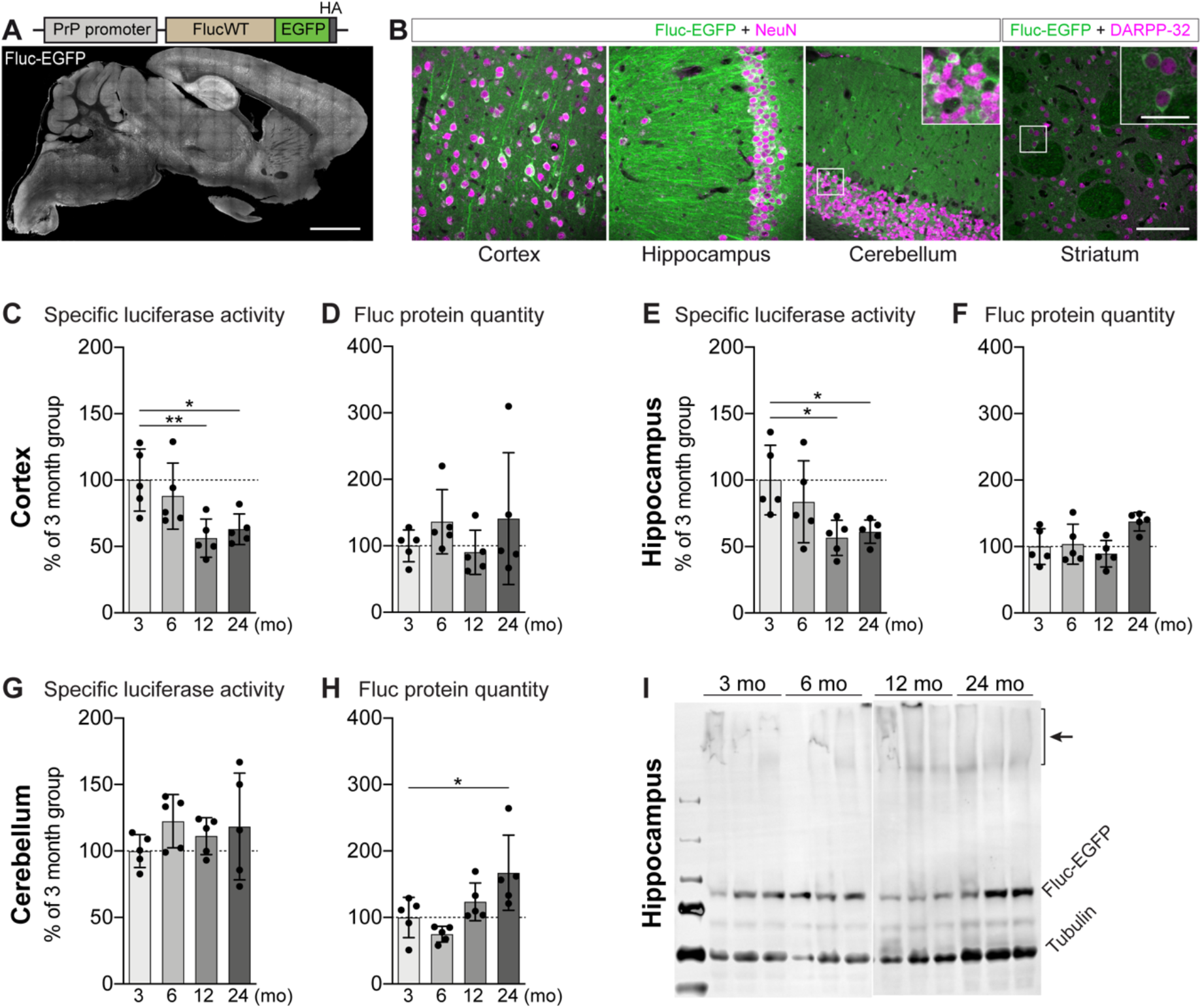
Fluc-EGFP reporter reveals proteostasis impairment in ageing mice. **A**, Scheme of the transgenic construct (top), and sagittal brain section of a Fluc-EGFP mouse from line 1214 at 3 months of age immunostained for EGFP (bottom). **B**, Representative images of the indicated brain regions of a Fluc-EGFP mouse, stained for EGFP (green) and the neuronal marker NeuN or the striatal medium spiny neuron marker DARPP-32 (magenta). Insets show higher magnification of the areas indicated by the boxes. **C-H**, Fluc-EGFP specific luciferase activity (C, E, G) and protein quantity (D, F, H) measured in the indicated brain regions of Fluc-EGFP mice at the indicated ages. Values are normalized to the 3- month-old group. N=5 Fluc-EGFP mice for each age group. One-way ANOVA with Bonferroni’s multiple comparisons test. ANOVA: C, **p = 0.0077; D, n.s; E, *p = 0.0205; F, *p = 0.0278; G, n.s; H, **p = 0.0058. Pairwise comparisons to the 3-month-old group are indicated on the graphs. **I**, Representative Western blot of hippocampal lysates at the indicated ages. Arrow points to the high molecular weight species observed in older mice. Several lanes on the blot between 6-month-old and 12-month-old samples were digitally removed. *p<0.05; **p<0.01. Scale bars: A, 2 mm; B, 100 μm; insets in B, 30 μm.

Unlike in cell culture conditions, the quantity of the Fluc-EGFP protein might vary considerably between tissue samples, which could lead to a bias when using bioluminescence measurements *in vivo.* We therefore first tested whether luciferase activity changes linearly to the Fluc-EGFP protein concentration using serial dilutions of 5 cortical tissue samples from 2-month-old Fluc-EGFP mice. We found that higher levels of Fluc-EGFP resulted in lower bioluminescence values than expected (Fig. S2A-B). The relationship between Fluc-EGFP protein quantity in the sample (x) and expected specific luciferase activity of the sample (y) was described by the formula: y = −0.45488x + 145.488 (Fig. S2C). All measurements of specific activity in tissue samples were therefore corrected accordingly.

Using Fluc-EGFP mice, we asked whether the reporter reacts to the *in vivo* changes in proteostasis capacity that are believed to occur in normal aging (Brehme et al., 2014; Klaips et al., 2018; Labbadia and Morimoto, 2015). To this end, we analyzed several brain regions of Fluc-EGFP mice at different ages (3, 6, 12 and 24 months). We observed a significant age-dependent decline in specific luciferase activity in the cortex and hippocampus, but not in the cerebellum (Fig. 2C-H). In 12- and 24-month-old mice, a high molecular weight smear was visible on the Western blot, likely corresponding to aggregated Fluc-EGFP species (Fig. 2I). These results show that Fluc-EGFP mice can be used to monitor *in vivo* alterations in proteostasis, and highlight intrinsic differences in the protein quality control of different brain regions.

### Fluc-EGFP sensor reacts to proteostasis defects in tauopathy mice

To investigate how proteostasis changes in disease, we crossed Fluc-EGFP mice to the rTg4510 line, a double-transgenic tauopathy model where expression of the responder transgene containing human four-repeat tau with the familial P301L mutation (tetO-tauP301L) is controlled by the CaMKIIα-tTA activator transgene. rTg4510 mice develop neurofibrillary tangle-like pathology and memory defects at 4 months, while neuronal loss and brain atrophy do not occur until 5.5 months of age (Santacruz et al., 2005). Consistent with previous reports (Myeku et al., 2016; Santacruz et al., 2005), immunostaining for phosphorylated tau (p-tau) revealed tau pathology in the cortex and hippocampus of 4-month-old rTg4510:Fluc-EGFP mice (Fig. 3A and S3A). At this time point, the majority of Fluc-EGFP positive cells contained multiple Fluc-EGFP foci (Fig. 3A-B and S3A). While foci were also observed in control mice, they were less frequent, and their fluorescence intensity was clearly lower than in rTg4510 mutants (Fig. 3A-D and S3A). Interestingly, the frequency of Fluc-EGFP foci in rTg4510 cortex was comparable in cells with and without p-tau (Fig. 3E), suggesting that Fluc-EGFP is sensitive to protein aggregation in the cytoplasm prior to formation of insoluble aggregates. Moreover, there was very little colocalization between p-tau and Fluc-EGFP foci in p-tau-positive cells (Fig. 3A, E and S3A), arguing that Fluc aggregation was truly due to impaired proteostasis rather than trapping of the sensor within the tau aggregates.

**Figure 3.**
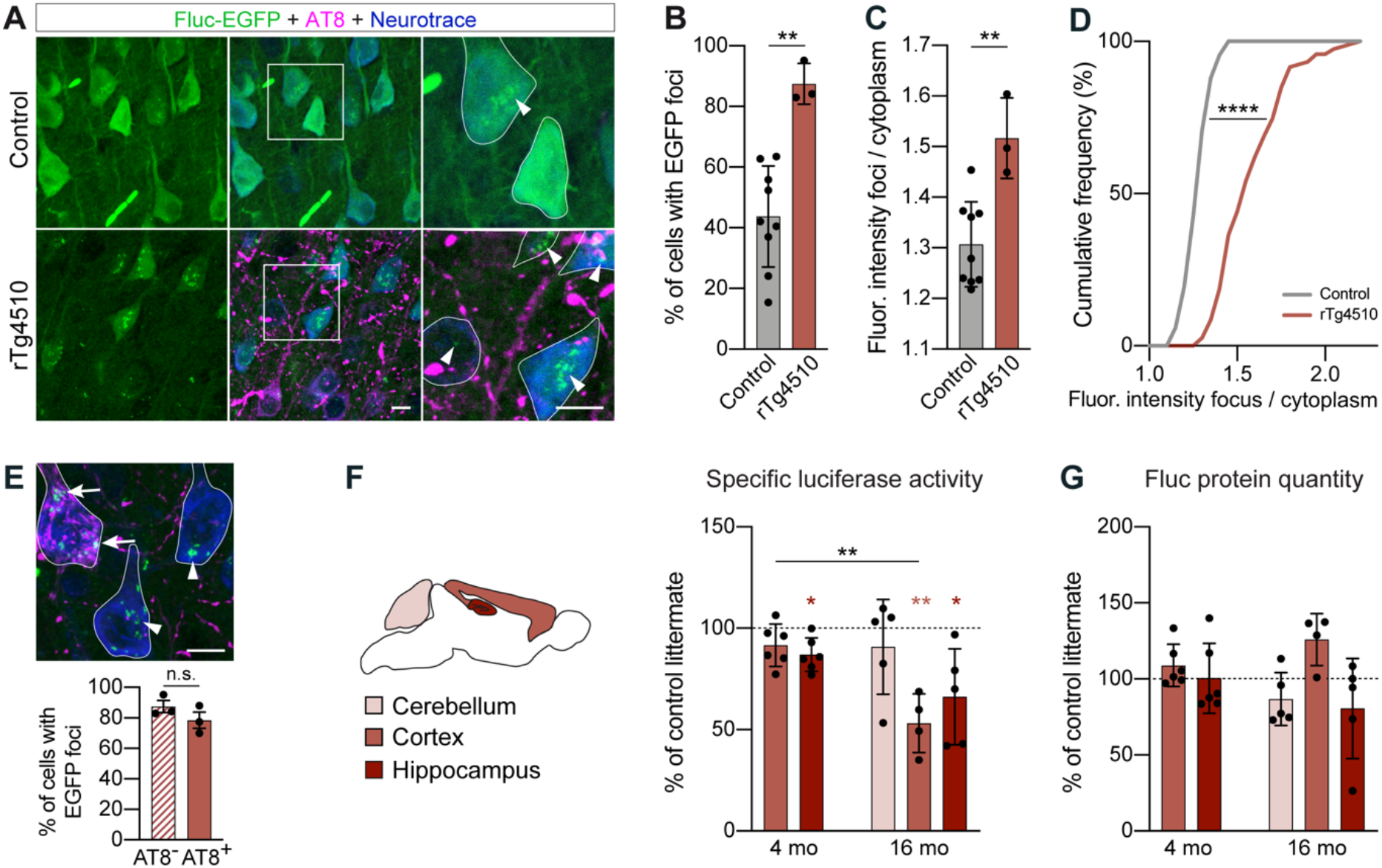
Fluc-EGFP reporter reveals proteostasis impairment in tauopathy mice. **A**, Cortical sections from rTg4510:Fluc-EGFP mice (lower row) and control littermates (upper row) stained for p-tau (AT8, magenta). Fluc-EGFP was detected by EGFP fluorescence (green), neurons were labeled with Neurotrace (blue). Images on the right show higher magnification of the areas indicated by the boxes, with neuronal cell bodies outlined. Note the presence of EGFP foci (arrowheads in the lower row) in the majority of neurons in rTg4510:Fluc-EGFP mice, and the absence of colocalization between EGFP foci and p-tau. Occasional foci of lower fluorescence intensity can be observed in control mice (arrowhead in the upper row). **B**, Quantification of the fraction of Fluc-EGFP positive cells with EGFP foci. N=3 rTg4510:Fluc-EGFP mice, 3 WT:Fluc-EGFP mice, 3 CaMKIIα-tTA:Fluc-EGFP mice and 3 tetO-tauP301L:Fluc-EGFP mice. WT:Fluc-EGFP, CaMKIIα-tTA:Fluc-EGFP and tetO-tauP301L:Fluc-EGFP littermates did not differ from each other and were pooled together as controls. Two-tailed t-test. **C**, Fluorescence intensity of EGFP foci, normalized to cytoplasmic EGFP fluorescence. Two-tailed t-test. **D**, Cumulative distribution of fluorescence intensities of EGFP foci. N=260 foci from 3 rTg4510:Fluc-EGFP mice and 364 foci from 9 control mice. Kolmogorov-Smirnov test. **E**, Example of EGFP foci in p-tau positive (arrows) and p-tau negative (arrowheads) cells (top), and quantification of the fraction of p-tau positive and negative cells containing EGFP foci (bottom) in rTg4510:Fluc-EGFP mice. N=3 mice. Two-tailed t-test. **F**, Left, scheme of brain regions used for luciferase assay color-coded by their vulnerability to tau pathology. Right, specific luciferase activity measured in the indicated brain regions of 4-month-old and 16-month-old rTg4510:Fluc-EGFP mice normalized to control littermates. N=6 rTg4510:Fluc-EGFP mice, 4 WT:Fluc-EGFP mice, 6 CaMKIIα-tTA:Fluc-EGFP mice and 1 tetO-tauP301L:Fluc-EGFP mouse at 4 months; 4-5 rTg4510:Fluc-EGFP mice, 5 WT:Fluc-EGFP mice, 3 CaMKIIα-tTA:Fluc-EGFP mice and 4 tetO-tauP301L:Fluc-EGFP mice at 16 months. WT:Fluc-EGFP, CaMKIIα-tTA:Fluc-EGFP and tetO-tauP301L:Fluc-EGFP littermates did not differ from each other and were pooled together as controls. Light-red and dark-red asterisks indicate comparisons to the corresponding littermate group (one-sample t-test), black asterisks indicate comparisons between corresponding brain regions of rTg4510:Fluc-EGFP mice at 4 vs. 16 months (two-tailed t-test). **G**, Fluc-EGFP protein quantity in the indicated brain regions of rTg4510:Fluc-EGFP mice measured by Western blot (for representative blots, see Fig. S3B). Values are normalized to control littermates. One-sample t-test, not significant for all groups. *p<0.05; **p<0.01; ****p<0.0001; n.s. - not significant. Scale bars in A and E, 10 μm.

The luciferase assay revealed a small, but significant decrease in specific Fluc enzymatic activity in the hippocampus, but not cortex of 4-month-old rTg4510 mice (Fig. 3F). The modest change in luciferase activity in total tissue lysates might be due to the variability of cellular reactions to misfolding *in vivo* at this early stage of pathology that might mask proteostasis impairments occurring in a fraction of cells. We therefore examined advanced-stage 16-month-old rTg4510 mice that show very abundant tau neurofibrillary tangles (Fig. S3A). At this age, a clear decrease in specific luciferase activity was detected both in the cortex and hippocampus, but not in the cerebellum, where the tau transgene is not expressed (Fig. 3F). Total Fluc-EGFP protein quantity was not significantly altered in any of the brain regions examined (Fig. 3G). However, high molecular weight species of Fluc-EGFP that were observed in all aged mice appeared more prominent in 16-month-old rTg4510 mutants, while the levels of Fluc-EGFP monomers were decreased compared to controls (Fig. S3B). These results suggest that the solubility of the Fluc-EGFP sensor decreases with the progression of tau pathology. Taken together, our findings in tauopathy mice demonstrate the ability of Fluc-EGFP to react to proteostasis defects caused by expression of an aggregating protein *in vivo.*

### Fluc-EGFP sensor does not detect proteostasis changes in HD mice

In addition to tauopathy mice, we investigated proteostasis in the R6/2 mouse model of HD. R6/2 is an early-onset model with a rapid disease progression, showing frequent mutant Huntingtin inclusion bodies (mHTT IBs) at the age of 3 weeks, and severe brain shrinkage along with marked motor impairments at 12 weeks (Burgold et al., 2019; Carter et al., 1999; Mangiarini et al., 1996; Meade et al., 2002). In contrast to rTg4510 mice, we did not observe any changes in cellular distribution of Fluc-EGFP in R6/2:Fluc-EGFP mice, even at the late symptomatic stage of 12 weeks when mHTT IBs were present in every cell (Fig. 4A). Consistently, no decrease in specific luciferase activity was observed in the cerebellum, hippocampus, cortex, or striatum of 12-week-old R6/2:Fluc-EGFP mutants compared to littermate WT:Fluc-EGFP controls (Fig. 4B), although measurements of protein quantity revealed a significant increase in Fluc-EGFP levels in the hippocampus, cortex and striatum of R6/2 mice (Fig. 4C). These results are compatible with a scenario where Fluc degradation might be impaired, but Fluc is nevertheless properly folded in the R6/2 brain.

**Figure 4.**
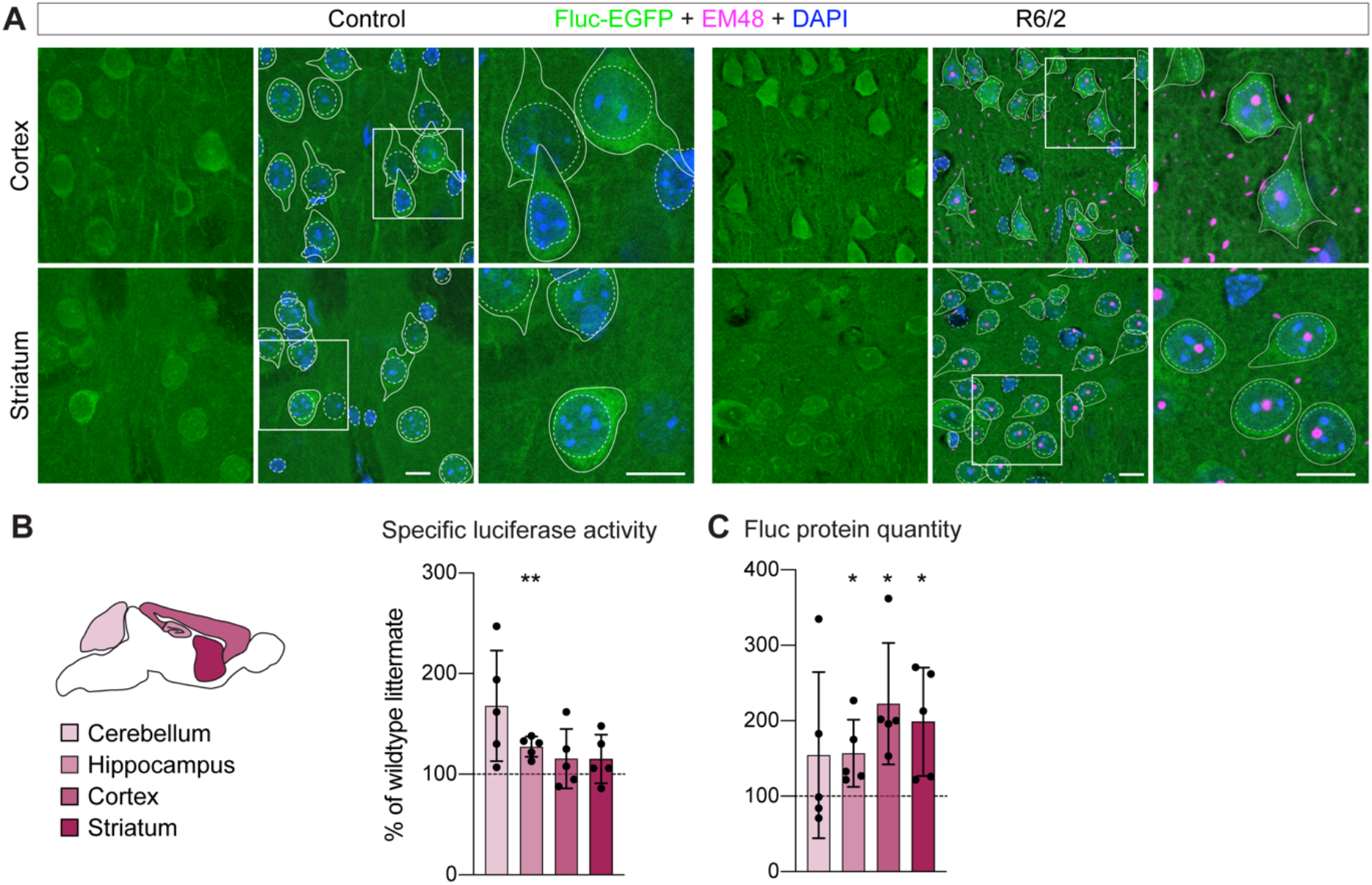
Fluc-EGFP reporter does not reveal proteostasis impairment in R6/2 mice. **A**, Cortical (upper row) and striatal (lower row) sections from 12-week-old R6/2:Fluc-EGFP mice (right) and control WT:Fluc-EGFP littermates (left) stained for aggregated mHTT (EM48, magenta). Fluc-EGFP was detected by EGFP fluorescence (green), nuclei were counterstained with DAPI (blue). Images on the right show higher magnification of the areas indicated by the boxes. Continuous and stippled lines mark cell bodies and nuclei, respectively. **B**, Left, scheme of brain regions used for luciferase assay color-coded by their vulnerability to Huntington’s disease. Right, specific luciferase activity measured in the indicated brain regions of 12-week-old R6/2:Fluc-EGFP mice normalized to WT:Fluc-EGFP littermates. N=5 mice of each genotype; One-sample t-test. **C**, Fluc-EGFP protein quantity in the indicated brain regions of R6/2:Fluc-EGFP mice measured by Western blot. Values are normalized to WT:Fluc-EGFP littermates. One-sample t-test. *p<0.05; **p<0.01. Scale bars in A, 10 μm.

Previous studies with UPS reporters also failed to show UPS impairments in HD mouse models with constitutive expression of mHTT (Bett et al., 2009; Maynard et al., 2009; Ortega et al., 2010). However, a transient defect in UPS was detected in the inducible HD94 model after mHTT expression was acutely switched on (Ortega et al., 2010). These previous observations raised the possibility that mHTT may initially impose a burden on the proteostasis system, which in the long term might be overcome by compensatory mechanisms. To test this, we crossed Fluc-EGFP mice to the inducible HD94 mouse line (CaMKIIα-tTA:BiTetO-HTT-Q94) (Yamamoto et al., 2000), which allows for precise temporal control over mHTT expression (Fig. S4A). The breeding pairs and the offspring were kept on Doxycycline throughout pregnancy and postnatally to suppress mHTT and prevent any compensatory adaptations (Fig. S4B). Doxycycline treatment was stopped at 8 weeks of age, and brains were harvested one and two weeks later, time points when expression of the mHTT transgene is already detectable and the most prominent UPS impairment occurs (Ortega et al., 2010). As mHTT is only expressed in the forebrain in the HD94 model, the cerebellum served as a negative control in these experiments. In agreement with our findings in the R6/2 line, no decrease in luciferase activity was observed in the hippocampus, cortex, or striatum of HD94:Fluc-EGFP mice 1 week or 2 weeks after mHTT transgene induction (Fig. S4C). We did not detect any differences in Fluc-EGFP protein quantity in this model compared to littermate controls (Fig. S4D). These data suggest that also acute expression of mHTT is not sufficient to cause proteostasis defects. Taken together, our findings indicate that the Fluc-EGFP sensor does not detect impairments of protein quality control in two different HD models.

### Cytoplasmic, but not nuclear, protein aggregates cause proteostasis impairments

One important difference between the disease models studied here is that in rTg4510 mice, aggregated tau is localized exclusively in the cytoplasm, including prominent pathology in the cell body, whereas both HD models display abundant mHTT IBs in the nucleus, but not in the cell body (Fig. 3A, E, S3A and 4A; Davies et al., 1997; Santacruz et al., 2005; Yamamoto et al., 2000). We therefore tested whether the Fluc-EGFP sensor is able to detect mHTT-dependent nuclear proteostasis impairments when targeted to the nucleus. To this end, we co-expressed FlucWT-EGFP versions targeted to the nucleus (NLS-Fluc-EGFP) and to the cytoplasm (NES-Fluc-EGFP) together with mCherry-fused versions of mHTT-exon1 that localize predominantly to the nucleus (mCherry-HTT-Q74, from here on nuc-mHTT) (Pan et al., 2018) or to the cytoplasm (HTT-Q97-mCherry, from here on cyt-mHTT) (Hipp et al., 2012). Both nuc-mHTT and cyt-mHTT readily aggregated and formed IBs in the respective compartment (Fig. 5A-B and S5B). Remarkably, cyt-mHTT caused proteostasis impairments that were detected by both cytoplasmic and nuclear targeted Fluc-EGFP (Fig. 5A-C). Conversely, nuc-mHTT did not cause foci formation neither by NES-Fluc-EGFP, nor by NLS-Fluc-EGFP (Fig. 5A-C). Moreover, in the presence of cyt-mHTT, we often observed mislocalization of NLS-Fluc-EGFP to the cytoplasm and appearance of cytoplasmic Fluc-EGFP foci, which were more abundant than nuclear ones (Fig. 5B, DE). Similar to our observations with p-tau in rTg4510 mice, no colocalization was detected between cyt-mHTT IBs and Fluc-EGFP foci (Fig. 5A-B). In summary, the Fluc-EGFP sensor reacts only to the presence of cytoplasmic, but not nuclear, protein aggregates, suggesting that only cytoplasmic protein aggregates cause a disturbance of neuronal proteostasis.

**Figure 5.**
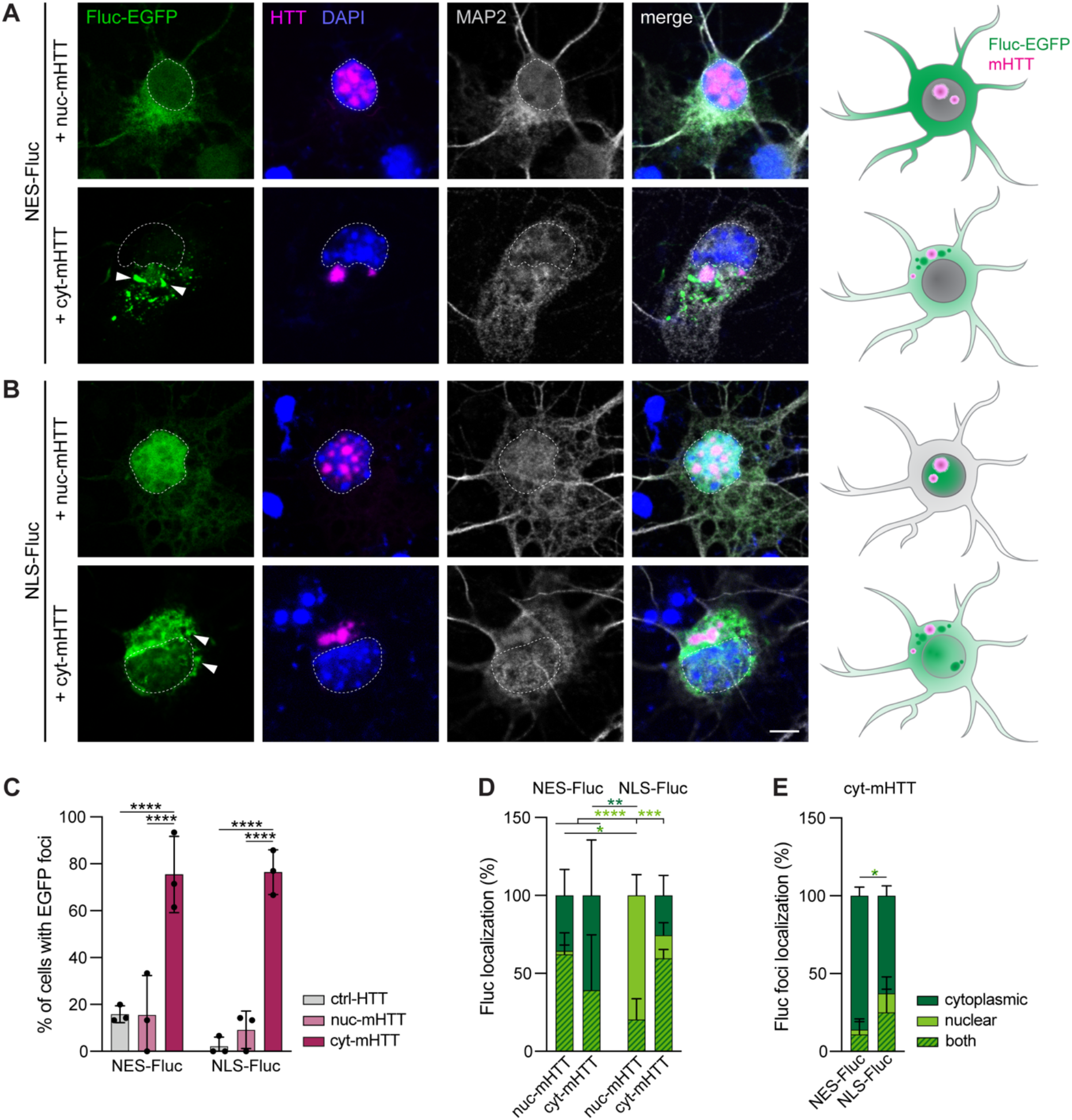
Response of NES-Fluc-EGFP and NLS-Fluc-EGFP to nuclear and cytoplasmic mHTT. **A-B**, Primary cortical neurons transfected with NES-Fluc-EGFP (A) or NLS-Fluc-EGFP (B) (green), in combination with nuc-mHTT (upper rows) or cyt-mHTT (lower rows) (magenta). Cells were fixed at DIV 3+2 and stained for the neuronal marker MAP2 (grey). Nuclei were labeled with DAPI (blue). Arrowheads point to Fluc-EGFP foci. Dashed lines mark the nuclei. Schemes on the right summarize the cellular distribution of the respective Fluc-EGFP (green) and mHTT-mCherry (magenta) proteins. Corresponding cultures transfected with control HTT are shown in Fig. S5A. **C**, Quantification of Fluc-EGFP foci formation. N=3 independent experiments. Two-way ANOVA with Tukey’s multiple comparisons test. ANOVA: HTT, ****p<0.0001; Fluc, n.s.; Interaction HTT × Fluc, n.s. Pairwise comparisons are indicated above the bars. **D**, Quantification of the subcellular localization of the respective Fluc-EGFP protein in the presence of the indicated mHTT constructs. N=3 independent experiments. Three-way ANOVA with Tukey’s multiple comparisons test. ANOVA: Localization, *p=0.019; HTT, n.s.; Fluc, n.s.; Localization × HTT, **p=0.0012; Localization × Fluc, ****p<0.0001; HTT × Fluc, n.s.; Localization × HTT × Fluc, ***p=0.0009. **E**, Quantification of Fluc-EGFP foci localization in the presence of cyt-mHTT. Two-way ANOVA with Sidak’s multiple comparisons test. ANOVA: Localization, ****p<0.0001; Fluc, n.s.; Localization × Fluc, *p=0.01. Dark-green, light-green and striped asterisks in D and E indicate comparisons of the fractions of cells with Fluc-EGFP localized in the cytoplasm, in the nucleus, or in both compartments, respectively. *p<0.05; **p<0.01; ***p<0.001; ****p<0.0001. Scale bar in B, 5 μm.

## Discussion

Here, we have generated new transgenic mice with expression of a proteostasis reporter in the brain, and demonstrated their utility in detecting proteostasis alterations in aging and disease. Proteostasis defects in the brain can be detected both by Fluc-EGFP foci formation, as well as by decline in luciferase activity. In contrast to available UPS reporters (Bove et al., 2006; Lindsten et al., 2003) that are limited to the analysis of protein degradation, Fluc-EGFP mice allow assessing the entire protein quality control system of neurons.

We used the Fluc-EGFP mice to probe proteostasis alterations in models of two different neurodegenerative proteinopathies, caused by expression of mutant tau and mHTT. In tauopathy mice, impairment of proteostasis was observed already at an early stage of disease. Of note, in our experiments with 4-month-old rTg4510 animals, impaired proteostasis could be detected more readily by changes in Fluc-EGFP cellular distribution than by changes in specific luciferase activity (Fig. 3A-D, F). This is in contrast to cell culture conditions where the luciferase activity readout showed a higher sensitivity to proteotoxic stress (Gupta et al., 2011). A possible reason for this difference might be the heterogeneity of cell types in the brain which differ in their proteostasis machinery and their responses to misfolding. These observations suggest that early changes in proteostasis might be difficult to detect with a luciferase activity-based assay in bulk tissue, and emphasize the importance of single-cell resolution approaches when studying protein misfolding in complex tissues *in vivo.*

In contrast to tauopathy mice, we found that proteostasis was largely intact in HD models. This is in line with previous studies using UPS reporters, which demonstrated UPS impairment in the rTg4510, but not R6/2 model (Bett et al., 2009; Maynard et al., 2009; Myeku et al., 2016). However, the observations we made in the HD94 line are seemingly in contrast to a previous report, where accumulation of the UPS reporter Ub-G76V-GFP was detected upon acute induction of mHTT expression (Ortega et al., 2010). Of note, our results do not exclude the possibility that protein degradation by UPS might be impaired in HD94 mice, however, they suggest that other components of the protein quality control machinery may compensate for the UPS defect.

We propose that subcellular localization of protein aggregates might be the cause of differences in proteostasis between tauopathy and HD mouse models. Indeed, we observed in cultured neurons that a mHTT fragment located in the cytoplasm had a profound effect on the solubility of the Fluc-EGFP sensor, while a mHTT fragment located in the nucleus did not. Our results therefore indicate that nuclear and cytoplasmic compartments differ in their capacity to cope with protein aggregation. Although mHTT is also found in the cytoplasm of HD mouse models used here (Landles et al., 2020; Osmand et al., 2016; Yamamoto et al., 2000), it does not seem to have a strong impact on the proteostasis network, in contrast to the effects of cytoplasmic mHTT IBs that we and others observed in cultured cells (Gupta et al., 2011). These differences might be caused by the localization of cytoplasmic IBs predominantly in neurites, but not in the cell body of neurons in HD mice, by higher expression levels of mHTT in cellular models compared to transgenic mice, and/or by the shorter duration of mHTT expression in cell culture, which provides less room for compensatory adaptations.

Previous studies in yeast and in cell lines described distinct protein quality control mechanisms operating in the nucleus and cytoplasm, and highlighted the unique role of the nucleus in maintaining cellular proteostasis and buffering the effects of aggregation (den Brave et al., 2020; Enam et al., 2018; Frottin et al., 2019; Park et al., 2013; Samant et al., 2018; Woerner et al., 2016). While our results are in agreement with these reports, we for the first time demonstrate the difference in proteostasis environment of the nucleus vs. cytoplasm in mammalian neurons *in vivo.* It should be noted that our findings do not exclude deleterious effects of nuclear mHTT, which have been demonstrated previously in cultured cells as well as animal models (Bae et al., 2006; Gu et al., 2015; Saudou et al., 1998; Veldman et al., 2015; Yang et al., 2002). Instead, they are compatible with the notion that nuclear and cytoplasmic mHTT aggregates have different molecular properties and are involved in different aspects of the disease (Landles et al., 2020). Our data suggests that toxicity of nuclear mHTT is likely due to other mechanisms than impaired proteostasis, and therefore might not be amenable to treatments targeting the proteostasis network. This provides a possible explanation for the weak effects of chaperone overexpression in mouse HD models that show prominent nuclear mHTT pathology (Hansson et al., 2003; Labbadia et al., 2012; Zourlidou et al., 2007), while similar strategies proved successful in rescuing toxicity caused by cytoplasmic polyQ fragments in cell lines and in flies (Gunawardena et al., 2003; Hageman et al., 2010; Jiang et al., 2012).

An unexpected finding in cyt-mHTT cells was the mislocalization of NLS-Fluc-EGFP to the cytoplasm (Fig. 5B, D). We speculate that it might be due to dysregulated nucleocytoplasmic transport that occurs in the presence of cytoplasmic aggregating proteins (Li and Lagier-Tourenne, 2018; Woerner et al., 2016), including mHTT (Gasset-Rosa et al., 2017; Grima et al., 2017).

In summary, our new proteostasis reporter mice represent a useful tool for detailed studies of proteostasis in various proteinopathy models, and can be used in the future for monitoring the success of therapeutic interventions that target the cellular protein quality control system. With the help of these mice, we uncovered unexpected differences between tauopathy and HD models that emphasize the unique proteostasis environment of the nucleus. Our results moreover raise the possibility that enhancing the proteostasis system as a treatment option might be more promising in diseases with cytoplasmic rather than nuclear localization of the aggregates. In future studies, it will be interesting to use the Fluc-EGFP reporter mice in the context of other protein misfolding diseases.

## Materials and methods

### Expression constructs

The following plasmids were used in the study: FlucWT-EGFP and FlucSM-EGFP (Gupta et al., 2011) with myc tag exchanged for HA; NLS-Fluc-EGFP and NES-Fluc-EGFP (Park et al., 2013); Mo.PrP (Borchelt et al., 1996); HTT-Q25-mCherry and HTT-Q97-mCherry (Hipp et al., 2012); EGFP-HTT-Q74 (Addgene 40262) with EGFP exchanged for mCherry. The sequences of the nuclear (mCherry-HTT-Q74) and cytoplasmic (HTT-Q97-mCherry, HTT-Q25-mCherry) HTT-exon1 constructs are similar apart from the polyQ length as well as the 7 N-terminal amino acids and 33 C-terminal amino acids of HTT-exon 1, which are present in the cytoplasmic, but not in the nuclear constructs.

### Mice

All the animal experiments were approved by the Government of Upper Bavaria (animal protocols 55.21-54-2532-13-13 and 55.2-1-54-2532-168-14) and conducted in accordance with the relevant guidelines and regulations. Mice were maintained in a specific pathogen-free animal facility with at libitum access to food and water. Animals of either sex were used for experiments.

For generation of Fluc-EGFP transgenic lines, FlucWT-EGFP and FlucSM-EGFP sequences with a C-terminal HA-tag were cloned into the Mo.PrP plasmid kindly provided by David Borchelt (University of Florida). Transgenic mice were generated by microinjection of linearized transgenic constructs into the pronucleus of C57BL/6 oocytes. Integration of the transgene was detected by PCR with the following primers: forward, 5′-GTG TCG CTC TGC CTC ATA GAA CTG CCT GCG TG −3′; reverse, 5′-CAT CCT TGT CAA TCA AGG CGT TGG TCG CTT CCG −3′. Fluc-EGFP mice were kept on C57BL/6 background.

R6/2 mice (Mangiarini et al., 1996) were obtained from JaxLabs (Stock No. 002810) and maintained by crossing transgenic males to F1 CBA/BL6 wildtype females. CAG repeat length was determined from tail biopsies by Laragen, Inc, and amounted to 192 ± 2 repeats. To obtain HD94 mice (Yamamoto et al., 2000), BiTetO mice (kind gift of José J. Lucas, Universidad Autonoma de Madrid) were crossed to the CaMKIIα-tTA line (Mayford et al., 1996) (JaxLabs, Stock No. 003010). HD94 breeding pairs and offspring were treated with 2 mg/ml Doxycycline, 5% sucrose in drinking water from conception until 2 months of age. rTg4510 mice (Santacruz et al., 2005) were obtained by crossing tetO-tauP301L mice (JaxLabs, Stock No. 015815) to the CaMKIIα-tTA line, and maintained on an FVB genetic background.

### Primary neuronal cultures

Sterile 13 mm glass coverslips were coated with 0.5 mg/ml poly-D-lysine overnight at 37°C and with 5 μg/ml Laminin in PBS for 2-4 hours at 37°C. Primary cortical neurons were prepared from E15.5 CD1 embryos. Pregnant females were sacrificed by cervical dislocation, the uterus was removed from the abdominal cavity, embryos were harvested and decapitated in ice-cold dissection medium consisting of Hanks’ balanced salt solution (HBSS) supplemented with 0.01 M HEPES, pH 7.4, 0.01 M MgSO4, and 1% Penicillin / Streptomycin. The skull was cut open and the cerebral hemispheres were separated from the rest of the brain. After removing the meninges, cortices were dissected and digested with prewarmed 0.25% Trypsin-EDTA supplemented with 0.75% DNAse for 15 min at 37°C. Trypsin activity was quenched by washing in Neurobasal medium containing 5% fetal bovine serum (FBS), and cells were dissociated in pre-warmed culture medium by triturating. Cells were centrifuged at 130 × g for 5 min and the pellet was resuspended in culture medium consisting of Neurobasal medium with 2% B27 (Invitrogen), 1% L-Glutamine (Invitrogen) and 1% Penicillin/Streptomycin (Invitrogen). Cells were plated in 24-well plates at a density of 100,000 per well. Transfection was performed at DIV 3 using CalPhos Mammalian Transfection Kit (TakaRa/Clontech). Briefly, coverslips were transferred into a new 24-well plate with fresh pre-warmed culture medium. Cells were incubated with 30 μl of the transfection mix (1.5 μg DNA, 124 mM CaCl_2_, 1 × HBS in H_2_O) at 37°C, 5% CO_2_ for 3 hours. A 24-well plate with fresh culture medium was equilibrated for at least 30 min in 10% CO_2_ and the coverslips were transferred into the equilibrated plate for 30 min at 37°C, 5% CO_2_. Finally, coverslips were moved back to the original medium and incubated at 37°C, 5% CO_2_ for protein expression. MG-132, Bafilomycin A1 and 17-AAG were diluted in 0.1% DMSO, and respective control samples were treated with 0.1% DMSO alone.

### Immunofluorescence

Mice were deeply anesthetized with 1.6% Ketamine / 0.08% Xylazine and transcardially perfused with PBS followed by 4% paraformaldehyde (PFA) in PBS. Brains were dissected out of the skull and postfixed in 4% PFA in PBS overnight. Fixed tissue was embedded in agarose and sectioned with a vibratome (VT1000S, Leica). Sections were re-hydrated in PBS and permeabilized with 0.5% Triton X100. After washing with PBS, cells were incubated in blocking solution consisting of 0.2% bovine serum albumin (BSA), 5% donkey serum, 0.2% lysine, 0.2% glycine in PBS for 30-60 min at room temperature (RT). Primary antibodies were applied in 0.3% Triton X-100, 2% BSA in PBS overnight at 4°C. The following primary antibodies were used: GFP (Life technologies, A11122), DARPP-32 (Lifespan Biosciences, LS-C150127), NeuN (Millipore, MAB377), AT8 (Invitrogen, MN1020), EM48 (Millipore, MAB5374). After washing three times with PBS, fluorescently labeled secondary antibodies and Neurotrace™ 435 / 455 (ThermoFisher, N21479, 1:200) were applied in 0.3% Triton X-100, 3% donkey serum for 1-2 hours at RT. DAPI was added at 1:1,000 dilution in PBS. Before mounting, the sections were treated with 0.5% Sudan Black B solution in 70% EtOH for 1 min in order to quench autofluorescence. Sections were mounted with fluorescent mounting medium (DAKO) or ProLong Glass Antifade medium (Invitrogen), and images were acquired with a Leica SP8 Confocal Microscope.

Cultured neurons were fixed with 4% PFA in PBS, washed with PBS, and permeabilized with 0.1% Triton X-100 in PBS for 10 min at RT. After washing with PBS, cells were incubated in blocking solution (2% bovine serum albumin, 4% donkey serum in PBS) for 30 min at RT. Primary antibodies were applied in blocking solution for 1 hour at RT. The following primary antibodies were used: cleaved Caspase-3 (Abcam, ab13847), MAP-2 (Novus, NB300-213). After washing three times with PBS, fluorescently labeled secondary antibodies were applied in blocking solution for 30 min at RT. DAPI was added at 1:1,000 dilution in PBS. Coverslips were mounted with fluorescent mounting medium (DAKO).

All images were acquired using a Leica TCS SP8 scanning confocal microscope. Images were processed and analyzed with Fiji-ImageJ. Quantification of localization and aggregation of Fluc-EGFP and HTT-exon1 in primary neurons was performed manually.

### Luciferase assay and Western blotting

Mice were sacrificed by cervical dislocation, brains were quickly removed from the skull, and brain regions were dissected on ice. Tissue samples were homogenized in pre-chilled lysis buffer (50 mM Tris-HCl, pH 7.4, 150 mM NaCl, 1% Triton X-100, protease inhibitor cocktail (Roche 04693132001) and phosphatase inhibitor (Roche, 04906837001)), centrifuged for 15 min at 14,000 × g and 4°C, supernatants were collected and used for the luciferase assay and Western blot. For luciferase assay, samples containing 100 μg total protein were mixed with 100 μl Luciferase Substrate (Promega). The kinetics of the luminescence was recorded for 10 min using a TriStar^2^ S LB 942 Plate Reader (Berthold Technologies) and the maxima of the curves were extracted. For Western blot, samples containing 100 μg total protein were denatured, separated on a 4-15% gradient gel (Criterion™ TGX Stain free™Precast Gel, Biorad), activated using a ChemiDoc Reader (Biorad) and transferred onto a PVDF membrane using a Trans-Blot Turbo transfer system (Bio-Rad). After blocking with 5% milk, 3% BSA, primary antibodies in 3% BSA, 0.1% Tris-buffered saline with 0.1% TWEEN® 20 (TBS-T) were incubated overnight at 4°C, followed by secondary antibodies in 3% milk, 0.1% TBS-T for 1 hour at RT. The following primary antibodies were used: GFP (JL-8, Clontech, #632381), Tubulin (Covance, MMS-435P). Proteins were visualized by fluorescence using the ChemiDoc Reader (Biorad). To take potential high-molecular weight smear of Fluc signal into account, the entire background-adjusted lane area above the Fluc-EGFP monomer band was quantified using Image Lab™ Software (Biorad).

### Data analysis

Statistical analyses were carried out with GraphPad Prism. Differences were considered statistically significant with p<0.05. Data are presented as mean ± standard deviation.

## Acknowledgements

We thank Frédéric Frottin for help at the initial stage of the project and for insightful discussions, Patricia Yuste-Chera for generously sharing reagents, David Borchelt for the Mo.PrP vector, José Lucas for the BiTetO mouse line, Tammo von Knoblauch, Raphaela Goetz, André Wilke, Sonja Schneider and Magdalena Böhm for support in mouse colony maintenance and genotyping, Julia Boshart for technical support with immunofluorescence, and Sonja Schneider and Patrick Auer for technical support with luciferase assays and Western blotting, respectively. This work was funded by the European Research Council (ERC) Synergy Grant under FP7 GA number ERC-2012-*SyG_318987*-Toxic Protein Aggregation in Neurodegeneration (ToPAG) (to F.U.H. and R.K.); and by the Max Planck Society for the Advancement of Science.

## Author contributions

M.S.H., F.U.H., R.K. and I.D. conceived the project. S.B., E.K.S.-T., R.K. and I.D. designed experiments. E.K.S.-T. performed proteotoxic stress experiments in cultured neurons, generated and characterized Fluc-EGFP transgenic mice, and conducted luciferase assays in R6/2 mice. S.B. and P.L. performed histology in young rTg4510 mice. S.B. and A.-L.B. performed histology in old rTg4510 mice and in R6/2 mice, and luciferase assays in aging Fluc-EGFP mice and in rTg4510 mice. I.D. and J.L. performed experiments with HD94 mice. A.-L.B. and K.V. conducted experiments with neuronal cultures co-transfected with Fluc-EGFP and mHTT. M.S.H. and F.U.H. provided tools and reagents. R.K. and I.D. supervised the project. S.B., E.K.S-T., K.V. and I.D. designed the figures. I.D. wrote the manuscript with input from all the authors.

## Conflict of interest

The authors declare no competing interests.

## Supplementary Figures

**Figure S1.**
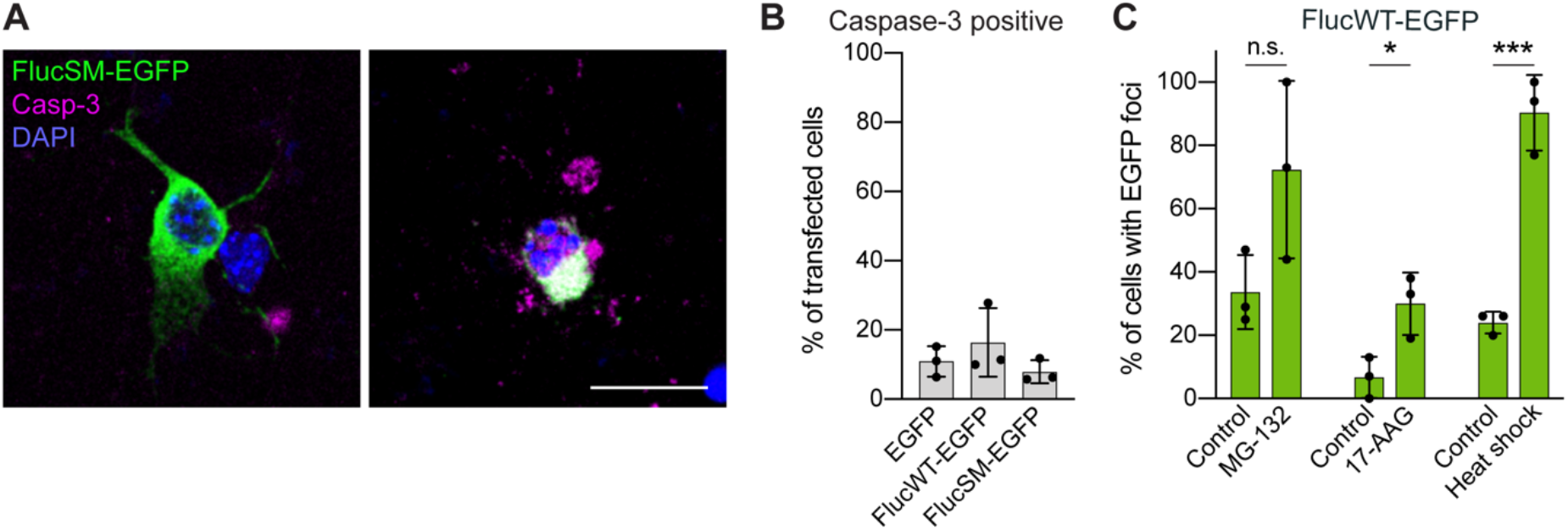
Fluc-EGFP reporters do not cause toxicity in primary neurons. **A**, Examples of neurons expressing FlucSM-EGFP (green) that are negative (left) or positive (right) for cleaved Caspase-3 (magenta). Nuclei are labeled with DAPI (blue). **B**, Quantification of the fraction of transfected neurons positive for cleaved Caspase-3 at DIV 3+2. N=3 independent experiments, 20 cells/condition/experiment; One-way ANOVA. No significant differences were observed. **C**, Quantification of FlucWT-EGFP foci formation in transfected neurons upon indicated treatments. N=3 independent experiments, 15 neurons/condition/experiment; Two-tailed t-test. *p<0.05; ***p<0.001; n.s. - not significant. Scale bar in A, 20 μm.

**Figure S2.**
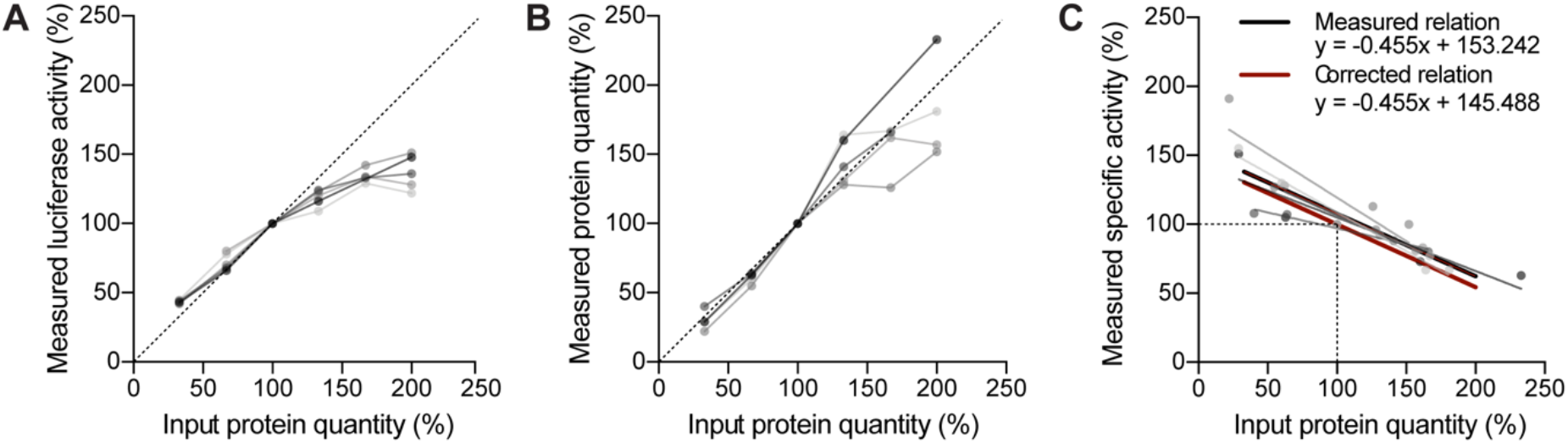
Linearity of luciferase activity in tissue samples. Cortical lysates from 5 Fluc-EGFP mice (shown in shades of grey) were used in different dilutions (25 μg, 50 μg, 75 μg, 100 μg, 125 μg and 150 μg input protein quantity). All values were normalized to the 75 μg dilution. **A**, Measured luciferase activity of the samples. **B**, Measured protein quantity. **C**, Linear regression of input protein quantity to specific luciferase activity (measured luciferase activity normalized to measured protein quantity) for individual mice (grey lines) and mean regression of five mice (black line); red line shows the corrected relation where 100% input protein quantity corresponds to 100% specific activity.

**Figure S3.**
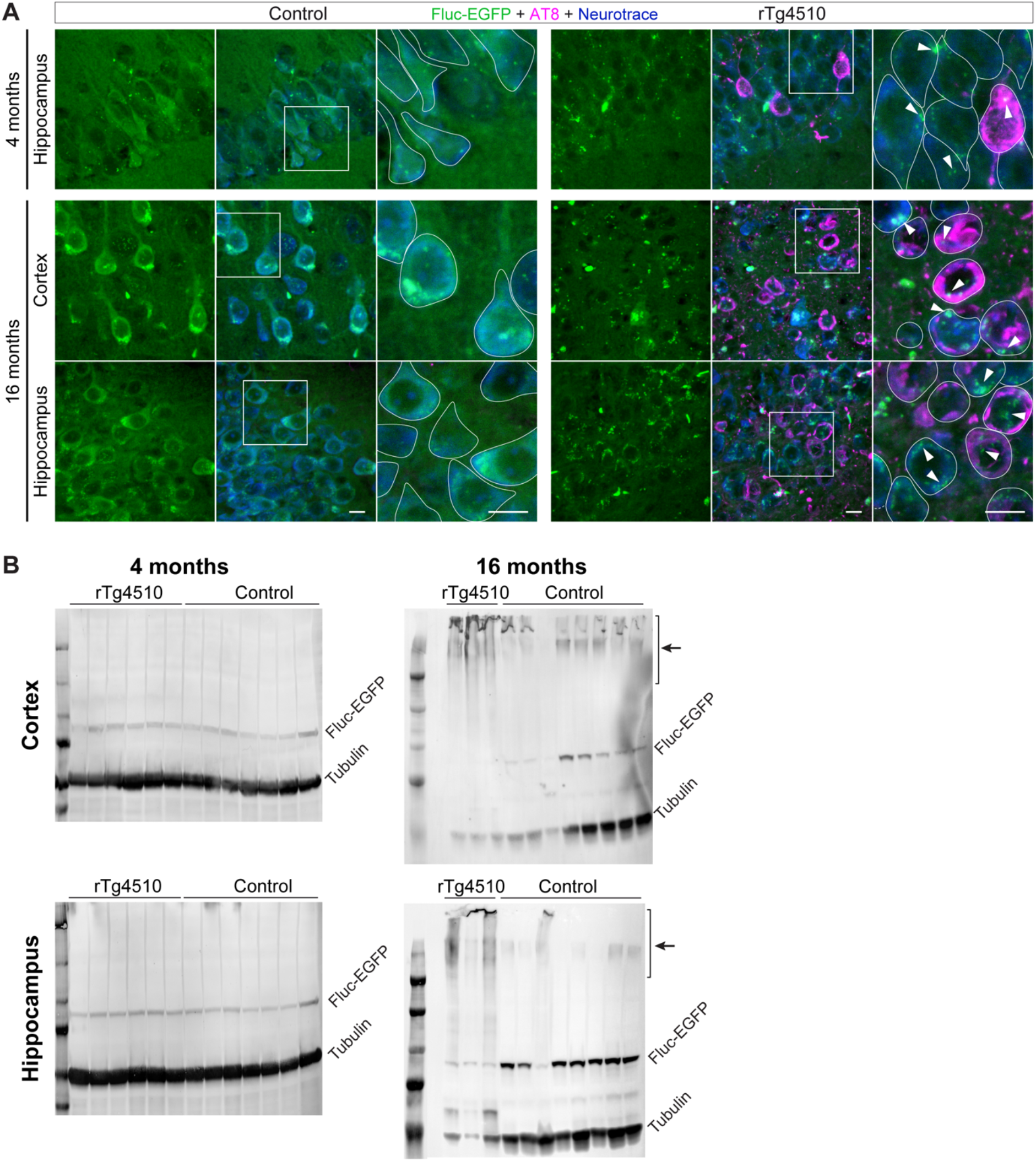
Additional analyses of Fluc-EGFP reporter in rTg4510 mice. **A**, Images of the cortex and hippocampus of control littermates (left) and rTg4510:Fluc-EGFP mice (right) stained for p-tau (AT8, magenta) at the indicated ages. Fluc-EGFP was detected by EGFP fluorescence (green), neurons were labeled with Neurotrace (blue). Higher-magnification images show the areas indicated by the boxes, with neuronal cell bodies outlined. Arrowheads point to Fluc-EGFP foci. **B**, Representative Western blots of cortical (top) and hippocampal (bottom) lysates from 4-month-old (left) and 16-month-old (right) rTg4510:Fluc-EGFP mice and control littermates. Arrows point to the high molecular weight species. Scale bars in A, 10 μm.

**Figure S4.**
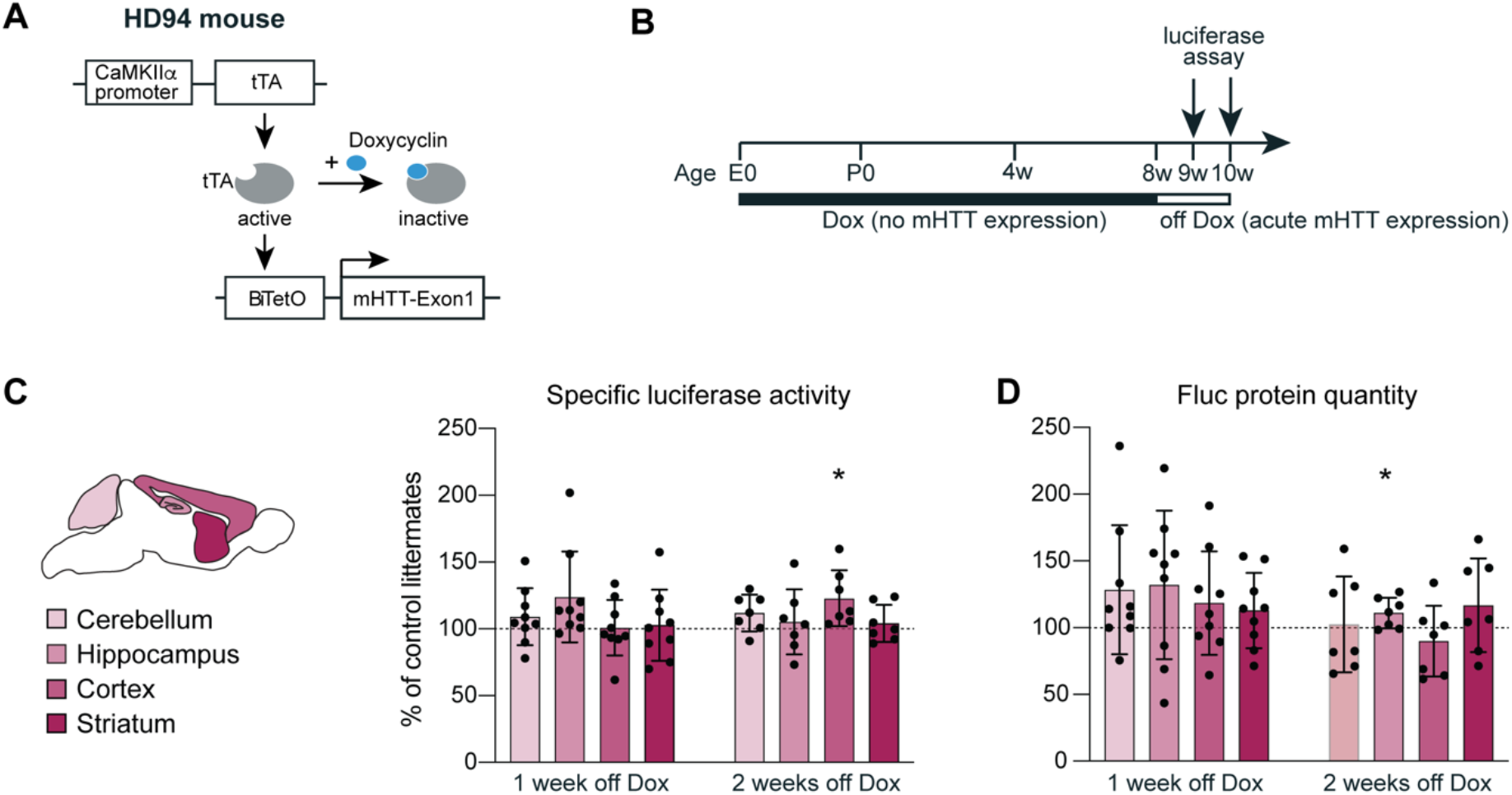
Proteostasis measurements in HD94:Fluc-EGFP mice upon acute induction of mHTT expression. **A**, Scheme of the HD94 transgenic strategy. **B**, Experimental timeline. E0, embryonic day 0; P0, postnatal day 0. **C**, Specific luciferase activity of cerebellar, hippocampal, cortical and striatal tissue lysates. **D**, Fluc-EGFP protein levels measured by Western blot. WT:Fluc-EGFP, CaMKIIα-tTA:Fluc-EGFP and BiTetO:Fluc-EGFP littermates did not differ from each other and were pooled together as controls. Values of HD94:Fluc-EGFP mice were normalized to their respective pooled littermate controls. 1 week off Dox: N=9 HD94:Fluc-EGFP mice; 5 CaMKIIα-tTA:Fluc-EGFP mice; 7 BiTetO:Fluc-EGFP mice; and 7 WT:Fluc-EGFP mice; 2 weeks off Dox: 7 HD94:Fluc-EGFP mice; 2 CaMKIIα-tTA:Fluc-EGFP mice; 3 BiTetO:Fluc-EGFP mice; and 3 WT:Fluc-EGFP mice. One-sample t-test, *p<0.05.

**Figure S5.**
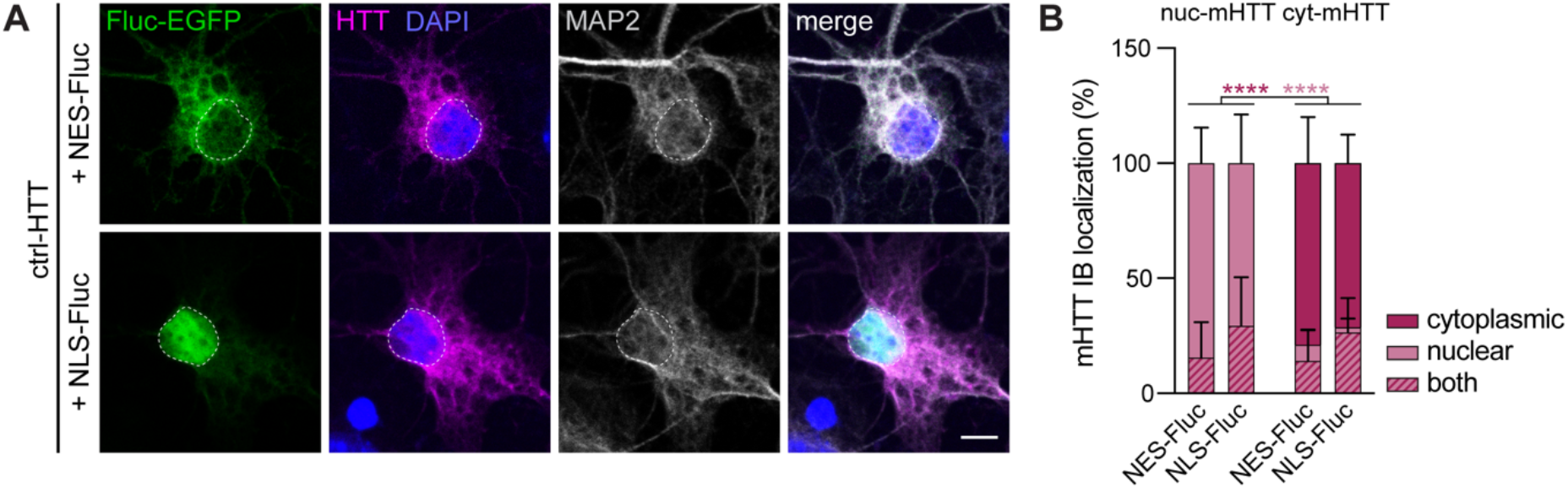
Co-expression of NES-Fluc-EGFP and NLS-Fluc-EGFP with control HTT, and analysis of subcellular localization of mHTT-mCherry variants. **A**, Primary cortical neurons transfected with NES-Fluc-EGFP (upper row) or NLS-Fluc-EGFP (lower row) (green), in combination with control HTT-exonl (HTT-Q25-mCherry, magenta). Cells were fixed at DIV 3+2 and stained for the neuronal marker MAP2 (grey). Nuclei were labeled with DAPI (blue). Dashed lines mark the nuclei. Corresponding cultures transfected with mHTT constructs are shown in Fig. 5A-B. **B**, Quantification of mHTT IB localization in the cytoplasm, nucleus, or both in the indicated conditions. Three-way ANOVA with Tukey’s multiple comparisons test. ANOVA: Localization, **p=0.0049; HTT, n.s.; Fluc, n.s.; Localization × HTT, ****p<0.0001; Localization × Fluc, n.s; HTT × Fluc, n.s.; Localization × HTT × Fluc, n.s. Dark-red and light-red asterisks indicate pairwise comparisons of the fractions of cells with mHTT IBs localized in the cytoplasm and in the nucleus, respectively. ****p<0.0001. Scale bar in A, 5 μm.

## Notes

### Competing Interest Statement

The authors have declared no competing interest.

